# Hippocampal Codes for Real-World Human Navigation

**DOI:** 10.1101/2023.06.22.545396

**Authors:** Kathryn N. Graves, Ariadne Letrou, Tyler E. Gray, Imran H. Quraishi, Nicholas B. Turk-Browne

## Abstract

As animals move through the world, the hippocampus represents their location, direction, and speed. Parallel studies in humans have been mostly limited to virtual navigation because of physical constraints of suitable neural recording technologies. However, there are known differences between real-world and virtual navigation, leaving open the question of how the hippocampus supports real-world navigation in humans. Here we report evidence from ambulatory patients with chronic brain implants that the location, direction, and speed of humans walking along a linear track are represented in local field potentials from the hippocampus. We further show in a subset of patients who were tested twice after long delays that these representations can be reliable over time. These findings provide the first demonstration of multiple, stable neural codes for real-world navigation in the human hippocampus.

---

The core behavioral ability of organisms to move through their environment and explore new places requires representations in the brain of real-world space and actions that can be performed in space. The discovery of place cells in the rodent hippocampus [1], with neuronal activity contingent on the animal’s position in its environment, was the first of many rodent studies to reveal specialized, navigation-related codes in the hippocampal formation. These include subsequent discoveries of grid cells in entorhinal cortex with place codes over multiple firing fields[2], time cells in hippocampus that code for specific time-points in sequential experience [3], head direction cells in postsubiculum that code for head direction in the horizontal plane [4], speed cells in the hippocampus that code for locomotion speed [5], boundary cells in subiculum that code for proximity to environmental boundaries [6], lap cells in hippocampus that code for number of laps around a continuous environment [7], and jump cells in the hippocampus modulated by gaps in navigated routes [8]. These codes exist at multiple levels of neural organization in rodents, from single units to local field potentials (LFPs) [9], and they combine to form an organism’s internal map of its environment.

Intracranial electroencephalography (iEEG) from epilepsy patients with acute brain implants for seizure monitoring provides a rare opportunity to test similar questions in humans. Place cells [10] and grid cells [11], as well as grid-like LFP activity [12], have been established in the human hippocampus and entorhinal cortex, respectively. However, unlike the large body of non-human animal work, these studies have relied on computer-based virtual navigation by necessity, to overcome the physical limitations of hospitalized iEEG patients. To date, only three published human studies have investigated hippocampal LFP during physical movement through space [13, 14, 15]. These seminal studies compared the neural signatures of movement versus non-movement under various conditions, for example revealing the modulation of hippocampal theta by the distance of self and others to environmental boundaries [15]. However, it remains unknown whether the human hippocampus represents many of the spatial codes observed in rodents during real-world navigation.

Beyond spatial coding in the moment, the stability of these representations across time is unknown in humans. Place coding in rodents and bats is often unstable over time: they can show global remapping of all measured place cells to new random field locations across navigation episodes [16] or partial remapping of some place cells to new fields while others maintain their fields [17]; though see [18]. In contrast, speed coding [5] and head direction coding [46] are more stable across navigation episodes. Whether and which hippocampal codes for real-world navigation are stable over time in humans has not previously been examined.

Which LFP frequencies represent features of human navigation is also unresolved. Neural activity in the theta band has been implicated in place coding in rodents [19], with theta phase procession, or the firing of place cells at earlier and earlier phases of the theta cycle, supporting spatial encoding in novel environments [20]. In humans, however, the analogous phenomenon during virtual navigation manifests in neural activity in the lower, delta frequency band [21]. Although this theta-delta difference was once attributed to a species difference between rodents and humans[22], it may instead reflect a difference between virtual and real-world navigation [13]. If so, hippocampal codes for real-world navigation may be represented in the theta band.

The objective of the current study was thus threefold — while recording hippocampal LFP from ambulatory human patients as they walked along a linear track, we sought to: (1) reveal hippocampal codes for location, direction, and speed during real-world navigation; (2) test the stability of these representations over time with repeated testing across days and weeks; and (3) identify the frequency bands of neural activity driving these effects.

## Results

### Spatial codes in human hippocampus

Twelve epilepsy patients (Table 1) implanted with electrodes [23] in one or both hippocampi were recruited for the study (Fig. 1a). Participants alternated between one-minute periods of walking and standing (cued by music) along a 32-foot linear track (Fig. 1b). Their position in space was recorded with a head tracker (Fig. 2a,b).

**Table 1.** Patient details

| Patient | Sex | Age | Model | Ch 1 | Ch 2 | Ch 3 | Ch 4 | Delay |
| --- | --- | --- | --- | --- | --- | --- | --- | --- |
| 1 | F | 31 | 320 | L Hipp | L Hipp | L ER | L ER | 21 |
| 2 | M | 54 | 320 | L Hipp | L Hipp | R Hipp | R Hipp | 84 |
| 3 | M | 65 | 320 | L Hipp | L Hipp | R Hipp | R Hipp | 16 |
| 4 | M | 36 | 320 | L Hipp | L Hipp | R Hipp | R Hipp |  |
| 5 | M | 29 | 300M | R Hipp | R Hipp | L Hipp | L Hipp |  |
| 6 | F | 60 | 320 | L Hipp | L Hipp | R Hipp | R Hipp | 4 |
| 7 | M | 53 | 320 | R Hipp | R Hipp | R LT | R LT |  |
| 8 | F | 66 | 320 | L Hipp | L Hipp | L AMT | L AMT |  |
| 9 | M | 64 | 320 | L Hipp | L Hipp | R Hipp | R Hipp |  |
| 10 | F | 47 | 320 | L Hipp | L Hipp | R Hipp | R Hipp |  |
| 11 | F | 33 | 320 | L Hipp | L Hipp | R Hipp | R Hipp |  |
| 12 | M | 39 | 320 | L Hipp | L Hipp | R SO | R SO |  |

Patient demographics (columns 2-3), RNS System model number (column 4), coverage of four channels (columns 5-8) and delay in days between Session 1 and Session 2 for four patients who participated twice (column 9). Hipp = hippocampus; ER = entorhinal; LT = lateral temporal; AMT = anterior mesial temporal; SO = suboccipital.

**Fig. 1.**
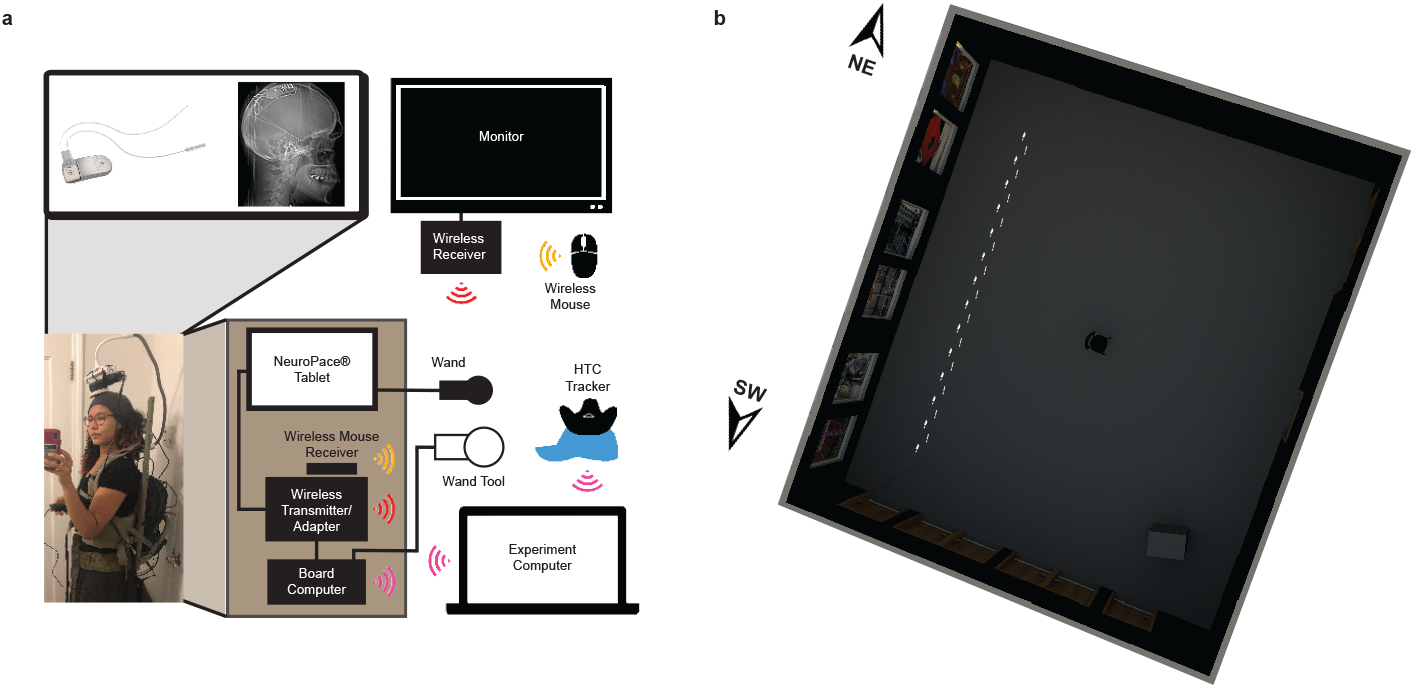
Data and environment. **a,** A telemetry wand and wand tool were affixed to the participant’s head (modeled in the photo by the first author of this manuscript) and connected to a clinical tablet and custom board computer continuously recording iEEG data. The tablet and board computer were stored in a backpack worn by the participant, and the tablet display was projected to a remote monitor via a wireless transmitter and receiver. A position tracker was placed on the participant’s head with Velcro on a felt cap. **b,** The study was performed in a 35 by 40 foot multimedia studio. Scene photographs were projected on screens along the long wall adjacent to 32 feet of walking space defined as the linear track.

**Fig. 2.**
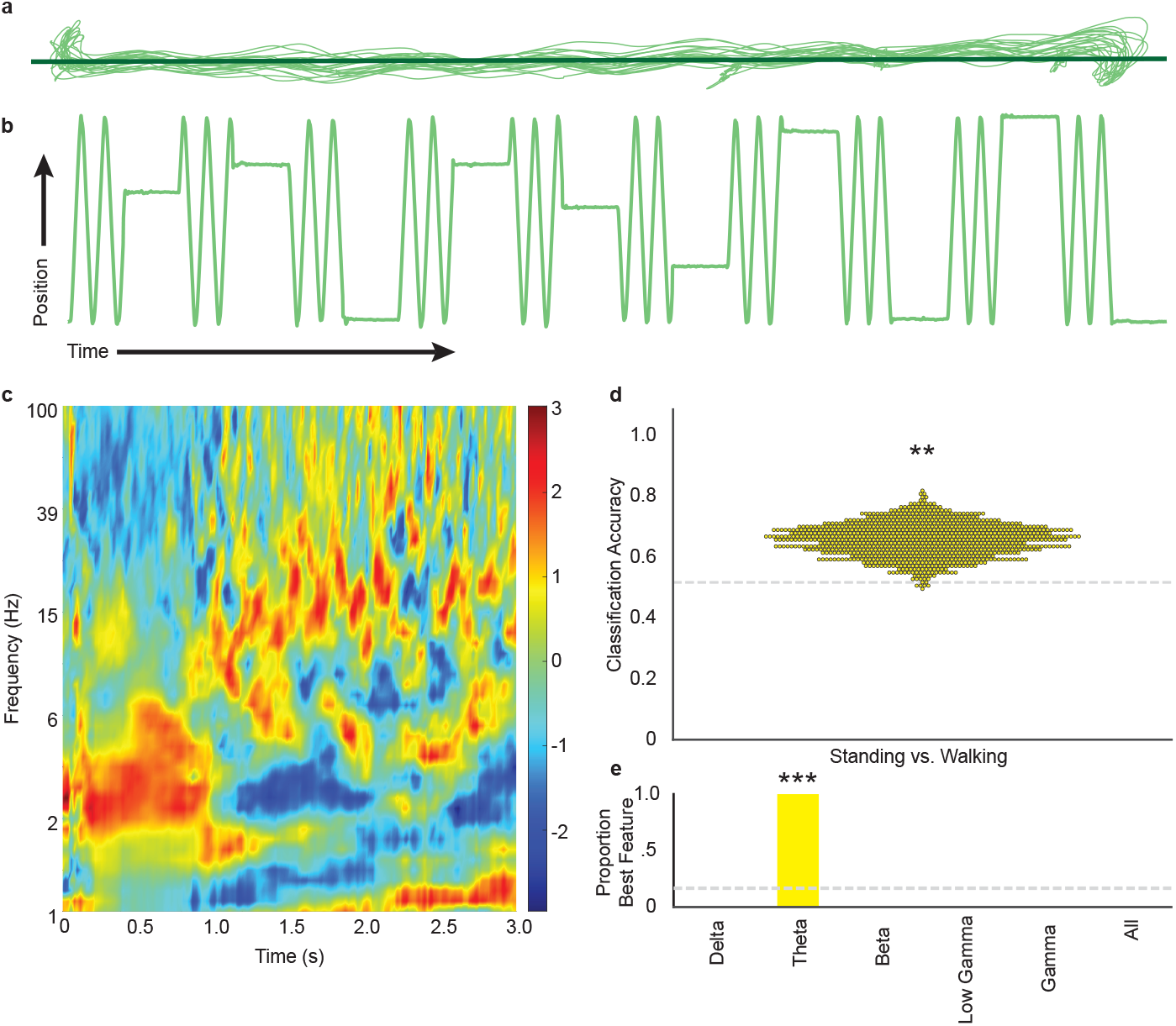
Walking and standing. **a,** Raw trajectories from bouts of walking in one example participant, overlaid with line of best fit calculated from the two-dimensional coordinates. **b,** Projecting these trajectories onto the line of best fit, plotted in transpose as the change in one-dimensional coordinates over time. Oscillations indicate bouts of walking back and forth. Plateaus indicate epochs of standing. **c,** Power from a time-frequency decomposition of hippocampal LFP for the onset and first three seconds of walking, averaged across all participants. **d,** Classification accuracy from decoding the onset of walking vs. standing, shown as bootstrapped sampling distribution at group level. **e,** Proportion of leave-one-participant-out (LOPO) cross-validation iterations in which the power at each frequency band yielded the highest classification accuracy. **P*<*0.01, ***P*<*0.001

### Movement

As an initial validation, we tested for known effects of movement on oscillatory activity in the hippocampus [38, 35]. We trained a classifier on a time-frequency spectral decomposition of hippocampal LFP data from all but one test participant at a time (Fig. 2c). The onset of walking vs. standing could be reliably decoded from the hippocampus of the held-out participant (Fig. 2d; boot-strapped P=0.008, 95% CI: [0.53, 0.74]). Leave-one-participant-out (LOPO) cross-validation was also used to determine which frequency band(s) were most informative (Delta: 1-4 Hz, Theta: 4-12 Hz, Beta: 12-30 Hz, Low Gamma: 30-60 Hz, Gamma: 60-100 Hz, All: 1-100 Hz). The theta band was selected on 100% of the iterations (Fig. 2e); this is more than would be expected by chance, defined as drawing from the six frequency bands at random (P<0.001).

### Direction

We next investigated spatial codes in the hippocampus during real-world navigation. The first type of spatial code we tested was heading direction: Southwest or Northeast (26°from true north/south). During periods of walking, Southwest vs. Northeast movement along the track could be decoded from spectral features of hippocampal LFP using LOPO cross-validation (Fig. 3a,e; bootstrapped P<0.001, 95% CI: [0.19, 0.95]). As a control, when standing and facing Northeast or Southwest, the direction could not be decoded reliably (P=0.823, 95% CI: [-0.69, 0.26]). Activity in the theta band drove decoding of walking direction on every iteration of LOPO cross-validation (Fig. 3b,c,d), more often than would be expected by chance (P<0.001).

**Fig. 3.**
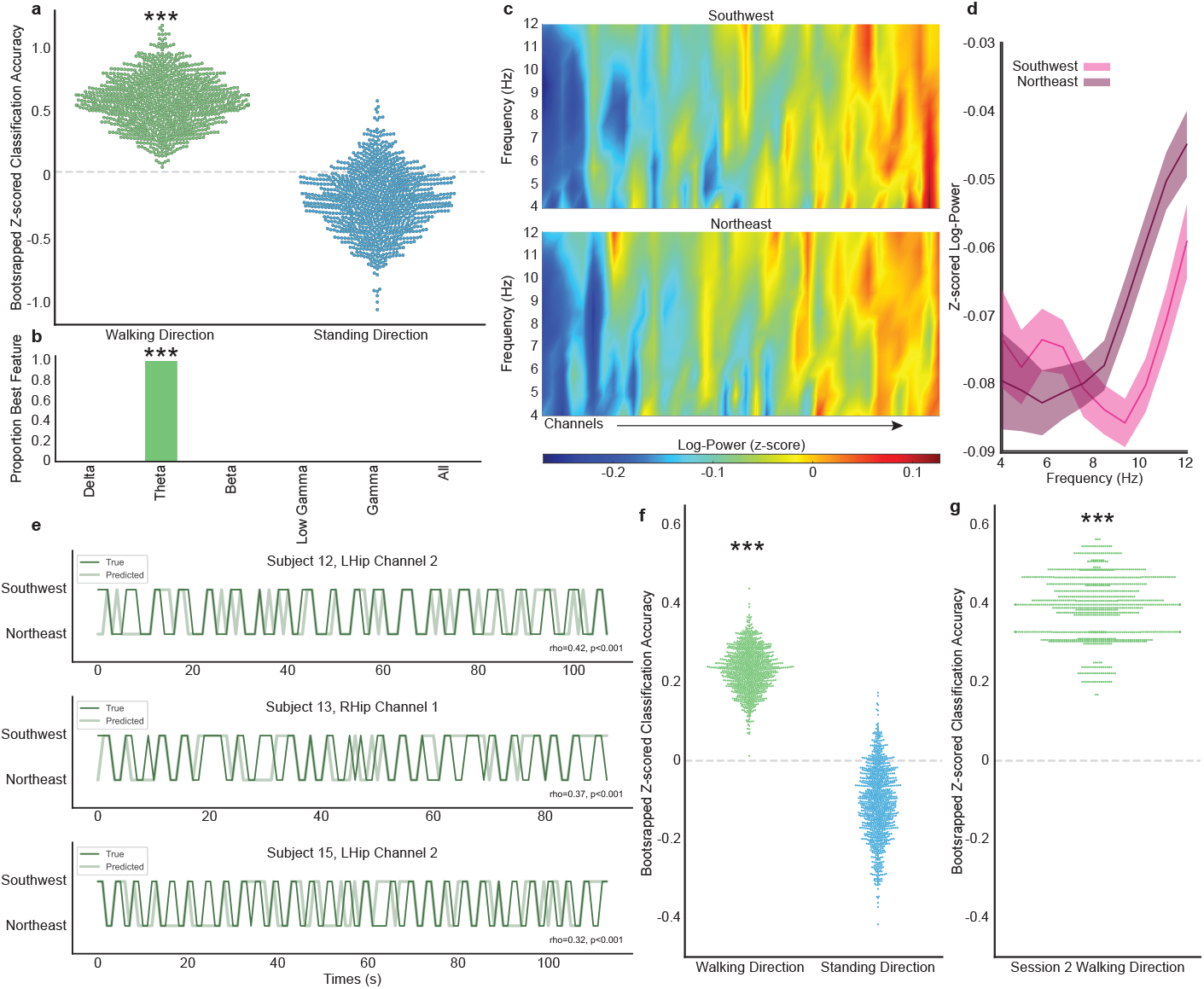
Heading direction. **a,** Classification accuracy from decoding the walking direction (green) and facing direction (blue) of a held-out test participant, shown as bootstrapped sampling distributions at group level. **b,** Proportion of LOPO cross-validation iterations in which the power at each frequency band yielded the highest classification accuracy for decoding walking direction. **c,** Average theta power spectra for each hippocampal channel pooling across participants, for Southwest (top panel) and Northeast (bottom panel) movement. Channels are ordered from lowest to highest average theta power during movement collapsed across direction and frequency. **d,** Power at each sampled frequency in the theta band during Southwest movement (light pink) and Northeast (dark pink) directions. **e,** Model predictions (light green) and true movement directions (dark green) over time for three example channels, downsampled for visibility. **f,** Alternative channel-level method in which classification accuracy was calculated separately for each channel, and channels rather than participants were resampled to determine significance. **g,** Cross-session classification accuracy of a model trained on Session 1 data and tested on Session 2 data; significant performance suggests that the neural features coding for direction were stable across sessions. ***P*<*0.001

### Location

We next tested for coding of location in the hippocampus during real-world navigation. Using regression and LOPO cross-validation, the current coordinate of a held-out participant while they walked on the linear track could be predicted from spectral features of windowed hippocampal LFP in the other participants (Fig. 4a,e; bootstrapped P<0.001, 95% CI: [0.26, 1.05]). Once again, activity in the theta band drove location prediction on every iteration of LOPO cross-validation (Fig. 4b,c,d), more often than would be expected by chance (P<0.001).

**Fig. 4.**
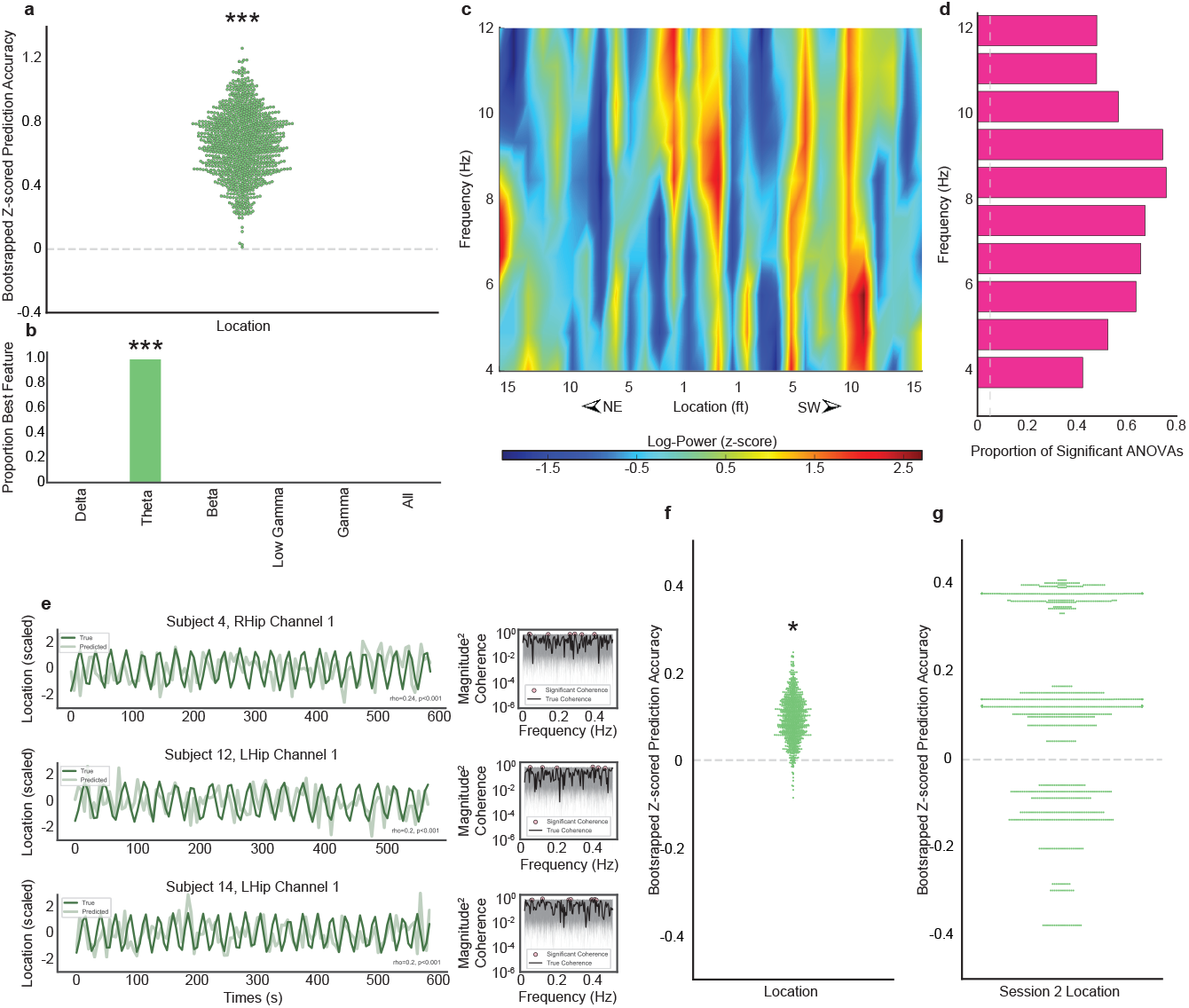
Track location. **a,** Regression accuracy in predicting the continuous location of a held-out participant on the track, shown as bootstrapped sampling distribution at group level. **b,** Proportion of LOPO cross-validation iterations in which the power at each frequency band yielded the highest accuracy in predicting spatial location. **c,** Average theta power spectrum across channels for coordinates along the linear track binned into 30 locations for visualization purposes. **d,** One-way ANOVAs quantified the relationship between track coordinates and hippocampal LFP power at individual theta frequencies. Average coordinates and LFP were defined by dividing the track into bins. The number of bins determined the granularity of the location coding. We considered a range of bin counts from 5 to *N*_coordinates_/2 bins in order to be robust to granularity. Across bin counts, the proportion of significant ANOVAs is plotted against the null alpha 0.05 significant tests by chance. **e,** Model predictions (light green) and true locations (dark green) over time for three example channels, downsampled for visibility. Inset plots depict significant coherence between true and predicted location trajectories. **f,** Alternative channel-level method in which regression accuracy was calculated separately for each channel, and channels rather than participants were resampled to determine significance. **g,** Cross-session prediction accuracy of a regression model trained on Session 1 data and tested on Session 2 data; chance performance suggests that the neural features coding for location can vary across sessions. *P*<*0.05, ***P*<*0.001

### Speed

Finally, we tested for coding of speed in the hippocampus during real-world navigation. Using regression, the current speed of a held-out participant while they walked on the linear track could be predicted from spectral features of windowed hippocampal LFP in the other participants (Fig. 5a,e; P<0.001, 95% CI: [3.00, 7.59]). Training the model simultaneously on all frequency bands from 1 to 100 Hz yielded the most accurate prediction of speed on nearly every iteration (92%) of LOPO cross-validation (Fig. 5b,c,d), more often than would be expected by chance (P<0.001); the remaining iterations (8%) selected the gamma band but this proportion was not reliable (P=0.89).

**Fig. 5.**
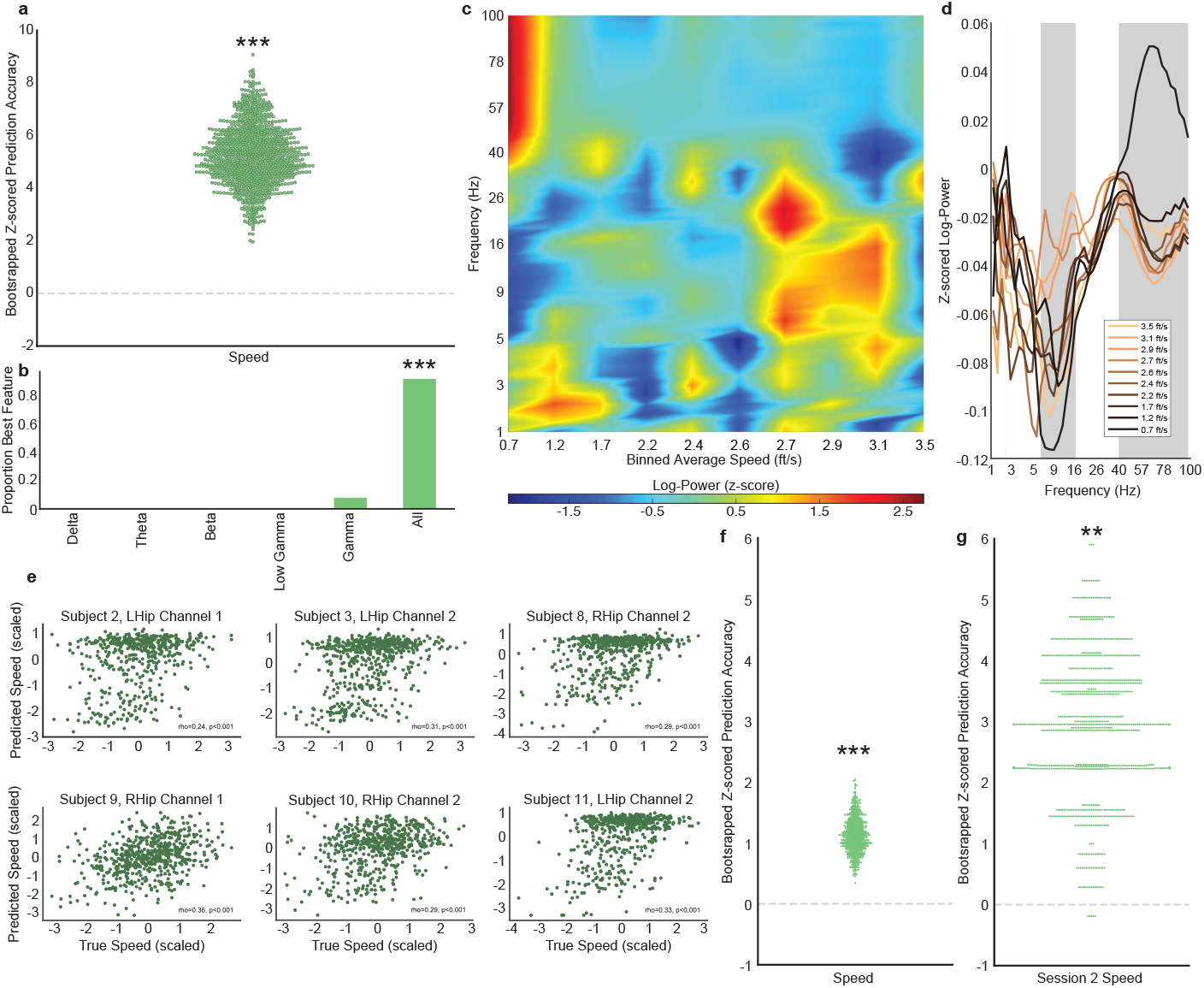
Walking speed. **a,** Regression accuracy in predicting the continuous speed of a held-out participant, shown as bootstrapped sampling distribution at group level. **b,** Proportion of LOPO cross-validation iterations in which the power at each frequency band yielded the highest accuracy in predicting speed. **c,** Average power spectrum across channels at different speeds, binned into 10 speed ranges for visualization purposes. **d,** Average power across channels at frequencies 1-100 Hz, binned into 10 speed ranges (separate lines) from slowest (dark brown) to fastest (light orange) speed. Gray shading indicates frequencies for which speed and power showed a significant Spearman rank correlation. **e,** Scatterplots of predicted and true speeds in example participants and channels. **f,** Alternative channel-level method in which regression accuracy was calculated separately for each channel, and channels rather than participants were resampled to determine significance. **g,** Cross-session prediction accuracy of a regression model trained on Session 1 data and tested on Session 2 data; significant performance suggests that the neural features coding for speed were stable across sessions. **P*<*0.01, ***P*<*0.001

### Representational overlap of spatial codes

We found evidence of spatial coding for heading direction, track location, and walking speed during real-world navigation when jointly considering all hippocampal channels for each participant (Supplementary Video 1a,b,c). Are these codes represented separately in distinct channels or do they overlap such that individual channels represent multiple codes? To address this question, we repeated the classification and regression analyses above within each of the 40 total hippocampal channels. We again found evidence, now at the level of individual channels, for coding of direction (Fig. 3f; walking: P<0.001, 95% CI: [0.13, 0.33]; standing: P=0.878, 95% CI: [-0.29, 0.06]), location (Fig. 4f; P=0.025, 95% CI: [0.001, 0.19]), and speed (Fig. 5f; P<0.001, 95% CI: [0.61, 1.71]).

Based on these results, we identified which channels showed above-chance performance for each code and counted the overlap of these channels across codes. Of the 40 total hippocampal channels, 28 coded for heading direction, 29 for track location, and 33 for walking speed. Of these, 21 coded for both direction and location, 23 coded for both direction and speed, 26 coded for both location and speed, and 19 coded for all three representations; these overlaps were all higher than expected by chance (Ps<0.001). Thus, a large proportion of hippocampal channels represented multiple spatial codes.

These overlapping spatial codes within a channel may be represented in different hippocampal units or dynamics that contribute to the population activity captured by LFP. We explored this possibility by testing whether direction, location, and/or speed were multiplexed in different frequencies. We quantified the overlap in the frequency bands selected during cross-validation and compared this to the overlap in a null distribution of randomly assigned frequencies. Speed was represented by different frequency bands than direction (P=0.02) and location (P=0.037), whereas direction and location did not diverge (P=1.0).

This overlap may also lead to interactions between spatial codes in the hippocampus. To examine this possibility, we performed a series of pairwise partial correlations by relating each channel’s evidence for one code (e.g., direction) with its evidence for another code (e.g., location), controlling for evidence of the third code (e.g., speed). There was no relationship across channels between the strength of direction and location coding (r=0.22, P=0.176, 95% CI: [-0.07, 0.52]) or location and speed coding (r=0.25, p=0.127, 95% CI: [-0.08, 0.56]). However, there was a significant *inverse* relationship between the strength of direction coding and speed coding (r=-0.37, p=0.022, 95% CI: [-0.61, -0.04]).

### Longitudinal stability of spatial codes

To explore the robustness and stability of spatial codes in the hippocampus over time, we were able to recruit a subset of four participants to return for a second experimental session (4–84 days after their first session; see Table 1). We repeated the channel-level analyses above for direction, location, and speed, first training the model on all of the hippocampal LFP data from Session 1 and then testing it on the data from Session 2. The direction classifier trained on hippocampal LFP from Session 1 could reliably decode heading direction during walking on Session 2 (Fig. 3g; P<0.001, 95% CI: [0.22, 0.53]). The speed regression trained on hippocampal LFP from Session 1 could likewise reliably predict walking speed (Fig. 5g; P=0.005, 95% CI: [0.60, 5.03]). However, the location regression trained on hippocampal LFP from Session 1 could not reliably predict location on Session 2 (Fig. 4g; P=0.283, 95% CI: [-0.30, 0.40]).

We further assessed the stability of these representations across time in the four repeat patients by quantifying the amount of overlap of above-chance channels on Session 1 and Session 2. There were 14 total hippocampal channels in these four patients. Of these channels, direction coding was observed in 13 channels in Session 1 and 8 channels in Session 2; all 8 channels overlapped across sessions, which was significantly greater overlap than expected by chance (p=0.010). Similarly, speed coding was observed in 12 channels in Session 1 and 13 channels in Session 2; 11 of these channels overlapped across sessions, which was significantly greater overlap than expected by chance (P<0.001).

## Discussion

In the current study, we demonstrated that the human hippocampus simultaneously represents the direction, location, and speed of a person freely navigating in the real world. Direction and location information were carried by spectral patterns in the theta band, whereas speed was best represented in broadband activity. Many individual recording channels represented more than one spatial code, suggesting that these codes are distributed and intermingled in the hippocampus. Finally, neural representations of direction and speed, but not location, were stable over a period of days to months in a subset of patients who were tested longitudinally.

These findings add to emerging evidence that neural population activity at the mesoscopic scale can mirror coding properties of single units [24]. For example, the spatial location of a rodent can be reconstructed from LFP with similar precision to spiking activity from individual neurons recorded simultaneously [9]. At a coarser scale, such representations have been observed in human brain regions [25]. Our study extends these findings to freely ambulating humans, demonstrating that place representations can be derived from hippocampal LFP across participants and from a single channel.

The finding that human theta encodes heading direction assigns a novel role to this established navigation-relevant frequency band. This complements previous studies of real-world human navigation that have linked hippocampal theta to fast vs. slow movement [13, 14] and to the proximity of environmental boundaries [15]. Importantly, coding of heading direction in hippocampal theta was limited to active walking and not to facing direction during standing. At a cellular level, head direction coding integrates landmark information and continuous changes in self-motion [36, 37]. Thus, heading direction may have only been apparent during walking because landmarks and self-motion were stable during standing.

The importance of theta in our findings and in extant real-world findings [13, 14, 15] contrasts with the dominance of the delta band during virtual human navigation [27]. This frequency difference may be driven by motor, proprioceptive, or other signals present in physical vs. simulated movement. However, it is important to acknowledge that the onboard recording filter settings (fixed for clinical purposes) may have attenuated activity in the delta band. Nevertheless, place coding in hippocampal theta in humans mirrors the animal literature [9], suggesting that prior observations of delta reflect a task rather than species difference.

In labeling which hippocampal channels carried information about direction, location, and speed, we discovered that many channels coded for more than one feature simultaneously. The recorded LFP might be comprised of activity from distinct neural populations, consistent with the finding that speed was multiplexed at different frequencies than direction and location within individual channels. Alternatively, a given neural population in the hippocampus may jointly represent multiple spatial (and non-spatial) variables [39]. Such neural reuse may result in competition between spatial representations, consistent with the observed negative correlation between speed and direction coding across channels. This mirrors recent rodent studies of an inverse relationship at the level of single units, but critically in parahippocampal cortex and not hippocampus [45].

The finding that direction and speed coding generalize longitudinally and in a consistent subset of channels suggests that the human hippocampus represents some navigational signals stably over time. This extends evidence from the rodent hippocampus that the preferred direction of head direction cells is stable across days [46] and that speed cells persist over time and testing environments [5].

Conversely, hippocampal representations of location along the track did not generalize over time. This could reflect remapping, which has been observed in up to 85% of recruited place cells in rodent hippocampus across sessions in repeated environments [44]. The extent of place cell remapping depends on several factors, including sensory conflict [17], hippocampal subregion [41], and individual differences [31]. These factors could be explored in future studies of channel remapping in human real-world navigation.

In sum, we uncovered population coding of spatial features in the human hippocampus essential for real-world navigation. These findings bridge human and non-human animal models of navigation and lay the foundation for relating the core navigational and mnemonic functions of human hippocampus. The ability to extract complex cognitive variables during naturalistic behavior from LFPs recorded with brain implants may also inform the development and applications of new brain-computer interface technologies.

## Methods

### Participants

Twelve patients (aged 29–65 years, mean=50 years, SD=7.1 years; 5 female) previously implanted with a neurostimulator with iEEG capability for treatment of medication-resistant epilepsy participated in the study for compensation. They provided informed consent according to a protocol approved by the Yale University Institutional Review Board. Placement of electrodes was determined on the basis of each patient’s clinical care.

### Task design

The study was conducted in a 35 by 40 foot multimedia studio (Fig. 1b). Participants performed a navigational task in which they alternated between epochs of walking along a linear track and standing still. The linear track extended 32 feet parallel to a longer wall of the studio with six vertical floor-to-ceiling windows covered by projection screens. We projected six images of unique outdoor landmarks onto these screens in a consistent order to encourage hippocampally driven landmark-based navigation [32]. We counterbalanced the spatial order of these landmarks on the screens across participants. Floor markers placed at the beginning and end of the walking area signaled where participants should turn around.

When to walk vs. stand was cued by one of two instrumental songs (each lasting 60 s) over a surround-sound system in the studio. Participants were instructed to walk back and forth along the linear track while the walking song played and to stand in place while the standing song played. Assignment of song to condition was counterbalanced across participants. The two songs were sampled from a free music database with Creative Commons licensing (freemusicarchive.org) and equated for beats per minute to within 1 beat using MIRtoolbox 1.7.2 [43]. There were 10 interleaved epochs of walking and standing, for a total of 20 epochs lasting 20 minutes, followed by a minute of standing in silence. There was a brief pause of 1 s between songs and an additional buffer of 10 s at the beginning and end of the task.

The position of the participant in the studio in X and Y coordinates was recorded in real-time at 120 Hz using an HTC Vive Tracker 2.0 (HTC Corporation) attached to the top of a felt hat using Velcro. The task script was coded in Python 3.5 to present songs and to query the Tracker using Pyopenvr python bindings for Valve OpenVR [33]. The script additionally inserted TTL pulses into the iEEG data at the onset and offset of each song with a custom-configured board computer, to synchronize the neural and behavioral data (see [34]).

### Data acquisition

The RNS System (NeuroPace, Inc.) is an FDA-approved chronic implant that provides closed-loop neurostimulation for the treatment of seizures. It consists of a generator with EEG acquisition, analysis, and stimulation circuitry, connected to up to two chronically implanted electrode leads placed at 1-2 seizure foci. The electrode locations are individualized based on clinical assessment of patients’ seizure onset location(s). Our setup (Fig. 1a) allows continuous wireless streaming and synchronization of iEEG data. Local field potential (LFP) data were acquired and continuously stored in 2 minute and 40 second segments via connection to a wand and wand tool using near-field telemetry.

This implanted device records electrophysiological data from up to 4 channels, each of which is a bipolar signal from neighboring electrode contact pairs. The sampling rate is 250 Hz, and signals are bandpass filtered in hardware with slightly different settings across models. To determine how much signal could be recovered at the low and high end of the spectrum, we conducted frequency response testing ex-vivo with the 300M and 320 models (see Table 1 for which patients had each model). Given our interest in frequency bands between 1 and 100 Hz, we assessed the amount of signal loss at each of these frequency limits: at 1 Hz, we could recover about 75% of the signal from the 300M and 80% from the 320; at 100 Hz, we could recover about 90% of the signal from the 300M and 95% from the 320. Given that we could recover signal at these limits despite the clinically required filters, we included in our analyses frequencies from the delta band (1-4 Hz) through to the gamma band (60-100 Hz). Furthermore, our results do not rely on comparing signal magnitude across bands, but rather on patterns of activity within band, which were not biased by the filter settings.

### Data preprocessing

Artifact rejection was performed manually for each channel by a trained reviewer, blinded to experimental condition. Timepoints containing epileptiform activity, stimulation, or telemetry loss were manually flagged for removal (5.63% of Session 1 data, 4.77% of Session 2 data). A 60-Hz notch filter was applied and the data was subsequently epoched into 1-minute trials. A Morlet wavelet decomposition was applied on each trial with 3 seconds of mirrored data added at the beginning and end to account for edge effects and subsequently discarded (wave number 5). The decomposition was conducted at 10 linearly spaced frequencies within each of the following bands: 1-4 Hz, 4-12 Hz, 12-30 Hz, 30-60 Hz, 60-100 Hz. All artifact-free segments of at least 3500ms in length were time-frequency decomposed. Power was computed, log-transformed, and z-scored. This z-scored power at linearly spaced frequencies within band served as the features in our classification and regression analyses. The participant’s position coordinates were interpolated to match the iEEG sampling rate. Instantaneous speed was calculated for each adjacent coordinate pair. Although participants were instructed to walk back and forth, their raw coordinates naturally deviated from a perfectly straight line. Thus, we collapsed the two-dimensional representation of position along the linear track to one dimension by first computing a line of best fit from their coordinates. We then snapped the raw coordinates to this line, rotated them such that all y-coordinates were set to zero, and shifted the x-coordinates such that the midpoint along the walked trajectory was 0 (Fig. 1a,b). Lastly, we coded which direction participants were moving in based on the sign of the difference between adjacent pairs of coordinates. To code in which direction they were facing during the standing epochs, we calculated the sign of the average difference between pairs of coordinates during the last second of the preceding walking phase.

### Data analysis

#### Classification of movement versus non-movement

We first aimed to replicate the classic finding of movement-modulated theta activity in the hippocampus. We extracted the first 3 seconds of each walking and standing epoch and averaged across time and epoch, resulting in a vector of average LFP power from 1 to 100 Hz per channel per participant for both walking and standing. Using an L2-penalized Linear Support Vector Classifier and LOPO cross-validation, we set aside one participant’s channels as a test set and used the remaining participants’ data to identify the best frequency of activity for decoding. In this inner loop, we assessed the decoding accuracy of each frequency using LOPO within the training set, fitting the model with the features of a given frequency and scoring it on the held-out inner loop participant. This gave us an average inner-loop accuracy per frequency, and the frequency yielding the highest accuracy was then used to fit the model tested on the outer-loop held-out participant.

After computing model accuracy for each participant, we calculated group confidence intervals using bootstrap resampling [42]. We sampled the classifier accuracy of the 12 participants with replacement 1,000 times and calculated the group average on each iteration to populate a sampling distribution. From this distribution, the 95% confidence interval was set as the 2.5th and 97.5th percentiles. Bootstrapped *p*-values were calculated as the proportion of the sampling iterations yielding average classifier accuracy lower than or equal to chance (.50).

We analyzed the contribution of different frequencies by tallying the number of times each band was selected during cross-validated model training. We used a binomial test to determine for which bands the count was above what would be expected by chance, given six frequency options (Delta, Theta, Beta, Low Gamma, Gamma, All) and therefore six possible outcomes.

#### Classification of direction

To decode in which direction the participant was facing, we first divided the data from each channel into 4-s subtrials of each direction. The number of subtrials per participant varied because of natural differences in walking speed. We averaged within each epoch over time, resulting in a vector of average LFP power from 1 to 100 Hz per epoch.

For the main group-level analyses, we assessed classification accuracy again using LOPO cross-validation. We iteratively held out all subtrials from all channels from one participant as the test set, used the remaining data from all other participants as the training set to select the optimal frequencies and train an L2-penalized Linear Support Vector Classifier via the same inner-loop model training procedure as above. We then tested this trained model on the held-out outer-loop data. Because the number of subtrials deviated from an exact 50/50 split across participants, we evaluated model performance for each iteration by shuffling the held-out test labels 1,000 times to produce a null distribution against which we calculated the *z* score of the true model performance. After computing this *z* -scored model accuracy for each participant, we again calculated group confidence intervals using a similar bootstrap resampling procedure. Bootstrapped *p*-values were calculated as the proportion of the sampling iterations yielding average accuracy with the opposite sign of the true effect. We again assessed the contribution of different frequencies using the same tallying and binomial test procedure.

For the channel-level analyses, we assessed classification accuracy by separately training and testing an L2-penalized Linear Support Vector Classifier on data from each channel using 5-fold cross-validation. For each fold, 20% of the data were held out as the test set and frequency selection and model training were performed on the training set. Within this inner loop, 20% of the training data were held out to evaluate the predictiveness of each frequency. Model accuracy was again quantified as a *z* -score relative to a null distribution. We calculated group confidence intervals using bootstrap resampling, here iteratively computing the group average accuracy from the channels of 12 patients sampled with replacement.

#### Prediction of speed and location

The steps for predicting participant speed and location were nearly identical to the classification pipeline above, with the key difference being the use of a Ordinary Least Squares Linear Regression for prediction of a continuous variable on the group-level, instead of classification of a binary label. On the channel-level, Linear Least Squares with L2 regularization, optimal when many features impact model performance, was used to continuously predict speed, given that changes in speed modulate broadband hippocampal power across frequencies in rodents [38, 30]. An L1-regularized Linear Model, optimal in cases of small numbers of significant parameters, was used to continuously predict location on the channel-level given that narrow-band power modulations encode rodent location [9].

#### Representational overlap

We evaluated overlap between spatial codes by testing whether the number of channels coding for more than one of direction, location, and speed was greater than would be expected by chance. Specifically, we computed the number of overlapping channels between each pair of two spatial codes and across all three. We then performed binomial tests to determine whether this overlap was greater than would be expected if each channel’s model accuracy was equally likely to be above or below chance for each code.

Given the significant overlap we identified, we then explored potential means by which multiple representations could simultaneously arise in individual channels – namely whether they emerged at different frequencies within a single channel, or interacted with other representations. To investigate the first possibility, for each channel, we quantified the number of frequency bands pulled out during each fold of model cross-validation that were common across each pair and all models (i.e. for one channel, if the frequencies selected one each of five folds were “delta, theta, theta, theta, beta”, for direction, “delta, theta, theta, theta, theta” for location, and “theta, beta, gamma, gamma, gamma” for speed, the overlap between direction and location would be three, the direction-speed overlap would be two, and the location-speed overlap would be one), and took the group average overlap per comparison. We then generated a null distribution of frequency overlaps by randomly sampling without replacement and assigning each channel five frequencies (emulating 5 folds) per representation, preserving the counts of each frequency from the true cross validation results (i.e. if a channel’s true cross-validation results were “delta, theta, theta, theta, theta”, two unique frequencies would be randomly sampled and one would be repeated four times). We did this 1,000 times to generate a distribution of overlap counts, and assessed significance as the number null average overlap counts that were fewer than the true overlap. We then searched for evidence of interactions between spatial codes. We performed Spearman’s partial-correlations between the model accuracies of each pair of representations, controlling for the third representation.

#### Longitudinal stability

To evaluate whether spatial codes were stable over time, we applied the classification of direction and prediction of location and speed models to a second session obtained with four participants. Namely, we trained the same models that were used for channel-level decoding on the data from Session 1, and tested them on the (held out) data from Session 2. Statistical significance of channel-level model performance was assessed using the methods previously described. The positions of the six projected images of unique outdoor landmarks were held constant across sessions.

Because direction and speed could be predicted reliably across sessions, we calculated whether the channels significantly representing these codes overlapped between sessions. For each of direction and speed, we performed a binomial test to determine whether the number of overlapping channels out of a total of 14 in these repeat participants was greater than would be expected if each channel’s model accuracy was equally likely to be above or below chance in either session.

## Supporting information

Supplemental Information Guide

## Data availability

Raw iEEG data from the RNS System may not be shared publicly due to restrictions from the manufacturer, however such data can be made available to other investigators upon reasonable request. We transformed these data into spectral power for the analyses and visualizations in this manuscript and will make those intermediate products publicly available in a permanent repository upon publication.

## Code availability

The custom computer scripts used to perform our analyses and generate the reported results will be made publicly available on GitHub upon publication.

## Acknowledgements

This work was supported by National Institutes of Health (NIH) grants F99NS125835 (to K.N.G.) and R01MH069456 (to N.B.T-B.), Swebilius Foundation grants (to I.H.Q and K.N.G.), and CTSA grant TL1TR001864 from the National Center for Advancing Translational Science (NCATS), components of the NIH, and NIH Roadmap for Medical Research (to K.N.G.). We thank Dr. Nanthia Suthana for her generous time and support in establishing the data collection apparatus; Drs. Joel Greenwood and Omer Mano in the Kavli Neurotechnology Core for developing custom hardware; Dana Karwas, Lauren Dubowski, and Ross Whitman in the Center for Collaborative Arts and Media, where we conducted the study; and the many patients and clinicians with whom we had the privilege of working.

## Author contributions

K.N.G.., I.H.Q, and N.B.T-B. designed the experiment. K.N.G. conducted the experiment and analyzed the data with support from A.L. and T.E.G. and feedback from I.H.Q and N.B.T-B. K.N.G. wrote the manuscript with feedback from all authors.

## Competing interests

The authors declare no competing interests.

## Notes

### Competing Interest Statement

The authors have declared no competing interest.

