## Supplemental Information Guide for "Hippocampal Codes for Real-World Human Navigation"

**Supplementary Videos 1a, 1b, 1c**
**File Titles:**
Supplementary_Video_1a.mov
Supplementary_Video_1b.mov
Supplementary_Video_1c.mov
**File Summary**: Animation of neural model predictions for real-world location, direction, and speed in three representative patients, reflected as footsteps and a target depicting true and predicted location, respectively, a compass reflecting predicted motion direction, and a bar depicting speed of movement.
**Title**: Neural model predictions of real-world location, direction, and speed in three patients
**Legend**: Real-world location, direction, and speed predicted from multivariate models of spectral features in local field potentials obtained from: (1a) Subject 4 right hippocampal channel 1, (1b) Subject 12 left hippocampal channel 1, and (1c) Subject 14 left hippocampal channel 1. Footsteps reflect the patient's true 1-dimensional trajectory, target reflects a regression model's continuous prediction at the current time point, with trailing footsteps and targets for the three previous locations and predictions. Compass arrow reflects a classifier model's binary prediction of heading direction (modeled over 4-s averaged epochs, interpolated to match the 1-s sampling rate of speed and location predictions). Bar plot reflects a regression model's continuous prediction of movement speed. Prediction time series were smoothed using a Gaussian kernel for visualization purposes.
